# Stress matters: a double-blind, randomized controlled trial on the effects of a multispecies probiotic on neurocognition

**DOI:** 10.1101/263673

**Authors:** Papalini S., Michels F., Kohn N., Wegman J., van Hemert S., Roelofs K., Arias-Vasquez A., Aarts E.

## Abstract

Probiotics are microorganisms that can provide health benefits when consumed. Recent animal studies have demonstrated that probiotics can reverse gut microbiome-related alterations in anxiety and depression-like symptoms, in hormonal responses to stress, and in cognition. However, in humans, the effects of probiotics on neurocognition remain poorly understood and a causal understanding of the gut-brain link in emotion and cognition is lacking. We aimed to fill this gap by studying the effects of a probiotics intervention versus placebo on neurocognition in healthy human volunteers.

We set out to investigate the effects of a multispecies probiotic (Ecologic^®^Barrier) on specific neurocognitive measures of emotion reactivity, emotion regulation, and cognitive control using fMRI. Critically, we also tested whether the use of probiotics can buffer against the detrimental effects of acute stress on working memory. In a double blind, randomized, placebo-controlled, between-subjects intervention study, 58 healthy participants were tested twice, once before and once after 28 days of intervention with probiotics or placebo.

Probiotics versus placebo did not affect emotion reactivity, emotion regulation, and cognitive control processes at brain or behavioral level, neither related self-report measures. However, relative to the placebo group, the probiotics group did show a significant stress-related increase in working memory performance after versus before supplementation (digit span backward, p=0.039, η^p2^=.07). Interestingly, this change was associated with intervention-related neural changes in frontal cortex during cognitive control in the probiotics group, but not in the placebo group. Overall, our results show that neurocognitive effects of supplementation with a multispecies probiotic in healthy women become visible under challenging (stress) situations. Probiotics buffered against the detrimental effects of stress in terms of cognition, especially in those individuals with probiotics-induced changes in frontal brain regions during cognitive control.

**Highlights:**

- We ran a randomized placebo-controlled fMRI study with a multispecies probiotic
- Probiotics did not affect neurocognitive measures of emotion and cognitive control
- Probiotics did affect stress-related working memory and neural correlates
- Probiotics in healthy individuals can support cognition under stress

## 1. Introduction

Probiotics are defined as bacteria providing health benefits to the host when consumed in adequate amounts (Hill et al., 2014). In the last few decades, an increasing number of animal studies have indicated a role of probiotics in regulating mood, emotional behavior, cognition, and response to stress (Cryan and Dinan, 2012; Smith et al., 2014). For instance, by means of probiotics it was possible to reduce anxiety-like behavior and to normalize brain-derived neurotropic factor (BDNF) in the hippocampus of mice with infectious colitis (Bercik et al., 2011), and to reverse the abnormal stress responses in germ-free mice (i.e. without a gut microbiome) (Sudo et al., 2004) and rats (Messaoudi et al., 2011). Other studies showed that probiotics are able to lower levels of systemic inflammatory cytokines (McCarthy et al., 2003) and to regulate central GABA receptor expression in mice (Bravo et al., 2011). Additionally, probiotics could normalize the immune response, as well as noradrenaline concentration in the brainstem of rats after maternal separation (Desbonnet et al., 2010). These beneficial effects of probiotics on mood and cognition are mediated by the bi-directional link between the brain and the gut microbiome, called the gut-brain axis (Cryan and Dinan, 2012).

In humans, six weeks of probiotic supplementation in patients with intestinal disorders was able to decrease the depressive complaints associated with the intestinal disease, which was related to decreased brain limbic reactivity to negative emotional stimuli (Pinto-Sanchez et al., 2017). In healthy humans, four weeks of fermented milk product supplementation was associated with decreased functional magnetic resonance imaging (fMRI) responses in affective, viscerosensory, and somatosensory brain regions during emotional face matching (Tillisch et al., 2013). Nonetheless, these latter results should be taken carefully due to a number of limitations, i.e. group sizes (ranging between 10-12 subjects) and probiotics’ effects versus the no-intervention group instead of versus the placebo group. Nonetheless, existing evidence seems to suggest effects of probiotics on neural emotion reactivity in humans. However, emotion reactivity is only one of different affective appraisal processes, which also consist of emotion-specific regulation and more generic cognitive control components (Etkin et al., 2015; Etkin et al., 2006; Kohn et al., 2014). Thus, although effects of probiotics have so far especially been observed on neural emotion reactivity, it is possible that this is related to regulation of emotion or to higher order cognitive control processes. Therefore, our first aim was to investigate whether emotion reactivity is specifically affected by probiotics, whether these effects occur through control of emotion (i.e. regulating automatic biases, see Etkin et al., 2015; Etkin et al., 2006), or can be seen independent of emotion, i.e. affecting cognitive control more generally (see e.g. (O’Hagan et al., 2017) in middle-aged rats).

We investigated the effects of a multispecies probiotics (Ecologic^®^Barrier) (Van Hemert, 2014), in a randomized, double-blind, placebo-controlled between-subjects design. This formulation has been tested before in both animal and human studies. Specifically, two rat experiments showed effects of this formulation on depressive-like behavior and on the transcript level of factors involved in HPA axis regulation (Abildgaard et al., 2017b; Abildgaard et al., 2017a). In a previous human study in n=40 healthy participants, 4-weeks supplementation with this product was associated with a reduction in self-reported cognitive reactivity to sad mood versus placebo (Steenbergen et al., 2015). We studied the effects of probiotics on neural correlates underlying emotion reactivity, its regulation, and general cognitive control, by using three robust cognitive paradigms during fMRI: the emotional face-matching task (Hariri et al., 2000), known to activate the limbic network, including the amygdala, involved in emotion reactivity; the emotional face-word Stroop task (Etkin et al., 2006) known to activate regions in ventral and dorsal medial frontal cortex involved in emotion regulation; and the color-word Stroop task (Stroop, 1953) known to activate frontal cortex regions involved in cognitive control (Cieslik et al., 2015).

Animal studies have also shown how the gut-brain axis is crucial for stress regulation, by influencing the development of the hypothalamic-pituitary-adrenal (HPA) axis, which – in turn – is related to mood, emotion, and BDNF expression important for learning and memory (Frohlich et al., 2016; Gareau et al., 2011; Li et al., 2009; Sudo et al., 2004). The effects of probiotics and stress on cognition might share common pathways of action (e. g. the HPA axis, Arnsten, 2015; Sarkar et al., 2016), however, it is unclear whether probiotics might affect cognitive performance independent or dependent of the detrimental effects of stress. Beneficial effects of probiotics under stress conditions have been clearly demonstrated in animal studies (Ait-Belgnaoui et al., 2014; Cowan et al., 2016; Messaoudi et al., 2011; Sudo et al., 2004), but probiotics’ effects on cognition and stress resilience in humans are scarce and sometimes contradictory (Allen et al., 2016; Kelly et al., 2017). Therefore, as secondary aim, we took into account the possibility that potential probiotics effects on cognition could exist as a consequence of an increased buffer against stress. The probiotic product under investigation, Ecologic^®^Barrier, is developed to strengthen epithelial barrier function and to decrease intestinal permeability for the endotoxin lipopolysaccharide (LPS), as demonstrated *in vitro* (Van Hemert, 2014). Human studies showed that acute-stress paradigms increase intestinal permeability to LPS (Alonso et al., 2012; Vanuytsel et al., 2014), and detrimentally affect memory performance in both rodents and humans (Gareau et al., 2011; Schoofs et al., 2009). For example, the socially evaluated cold pressor test (SECPT) (Lovallo, 1975) specifically influenced backwards digit span (DS) performance, which involves control functions to operate on the stored material instead of just working memory maintenance (Schoofs et al., 2009). For this reason, we investigated whether the use of probiotics can modulate working memory (i.e. backwards DS) performance before versus after acute stress induced by the SECPT, together with stress-related changes in hormones (i.e. cortisol and alpha-amylase) and cardiovascular activity. As the type of cognition we investigated after and before stress – i.e. backwards digit span - requires cognitive control (Kane and Engle, 2003), we also investigated how intervention-induced effects on this stress-related working memory performance related to the effects of the intervention on cognitive control responses in the frontal cortex.

## 2. Methods

### 2.1 Participants

In total, fifty-eight of the 60 scanned participants were included in the analyses, divided into a probiotics intervention group (n = 29, mean age = 21 years, SEM = 0.4) and a placebo group (n = 29, mean age = 22 years, SEM = 0.5). Two participants were excluded from the analyses, one due to high depression levels (above BDI cut off for moderate depression, i.e. BDI score: 23), and one dropped out after the supplementation period. All participants were right handed, healthy female volunteers aged between 18 and 40 years old, using (oral or intra-uterine) hormonal contraceptives, with a healthy weight, i.e. a body mass index (BMI) between 18 and 25 (placebo group: mean BMI = 21.66 kg/m^2^, SEM = 0.31, and probiotics group: BMI = 21.91 kg/m^2^, SEM = 0.29). They were not in the ‘stop week’ of oral contraceptives during test sessions to ensure similar hormone levels between both sessions across participants. Exclusion criteria included: 1) personal history of psychiatric, neurological, gastrointestinal, endocrine disorders, and relevant medical history (self-reported); 2) regular medication use; 3) pre- and probiotic supplementation; 4) smoking; 5) use of antibiotics within two months before the start of the study. We also excluded those participants following a vegan diet, and those with high alcohol intake (i.e. more than 10 glasses per week). Furthermore, participants were screened for MRI compatibility. In order to ensure good task comprehension and clear understanding of the neuropsychological questionnaires, all participants exhibited sufficient knowledge of Dutch. The study was conducted following the Declaration of Helsinki with human subjects and the complete procedure was approved by the local Ethics Committee (CMO Arnhem-Nijmegen, NL55406.091.15) and registered at the Dutch trial register (protocol number: NTR5845). Written informed consent was obtained from each participant.

### 2.2 Intervention

Probiotics and placebo were consumed in powder form, 2 grams once daily at a fixed time point, on an empty stomach by diluting the powder in water or milk. Participants were asked not to eat for the subsequent 15-20 minutes after the ingestion of the drink. Ecologic^®^barrier consisted of the following bacterial strains: *Bifidobacterium bifidum* W23, *Bifidobacterium lactis* W51, *Bifidobacterium lactis* W52, *Lactobacillus acidophilus* W37, *Lactobacillus brevis* W63, *Lactobacillus casei* W56, *Lactobacillus salivarius* W24, *Lactococcus lactis* W19 and, *Lactococcus lactis* W58. With the application of new molecular identification techniques (including whole genome sequencing), the declaration of bacterial strains has been updated compared to previous publications (Abildgaard et al., 2017b; Abildgaard et al., 2017a)(Steenbergen et al., 2015). It has been confirmed that the probiotic formulation has always contained these nine strains, and has not been changed in ratio or CFU count since it has been researched. The strains (total cell count of 2.5 x 10^9^ colony forming units –cfu- per gram, i.e. 5 x 10^9^ cfu per day) were blended into a carrier material consisting of maize starch, maltodextrin, vegetable protein and a mineral mix. The placebo consisted of the same carrier material as used in Ecologic^®^Barrier and was indistinguishable in color, smell, taste and appearance.

All participants were randomly assigned to the probiotics or the placebo group. The randomization scheme was computer generated by Winclove using permuted blocks with block size equal to 4. It was impossible for research personnel involved with participants to adjust randomization or discern what product participants were receiving, ensuring true allocation concealment.

### 2.3 Procedure

#### 2.3.1 General procedure

A longitudinal double-blind randomized design was used to compare the effects of probiotics with placebo. Each participant was assessed twice: before the start of the treatment and four weeks later. Between the test sessions, a 28 days intervention consisting of probiotics or placebo intake was implemented. Both test sessions were conducted at the Donders Centre for Cognitive Neuroimaging in Nijmegen, The Netherlands. At the beginning of the first test session, the experimental procedures were explained and the principal researcher assessed physical measurements, including height, weight, blood pressure, and heart rate (Figure 1). Next, participants practiced all the fMRI tasks outside the scanner, and performed a working memory test (i.e. backward and forward digit span test). Subsequently, the participants took part in the fMRI experiment (75 minutes), with three cognitive paradigms described below. Within 10-15 minutes after the MRI measurement, another trained researcher (unfamiliar to the subject) conducted a stress task followed by the same working memory test (with different items) as performed before scanning (conducted by the principal researcher again). Saliva and cardiovascular parameters were collected over the testing sessions (see Figure 1, for details see Supplementary Materials). At the end of the first session day, participants were provided with the probiotics/placebo (in identical sachets, blind to both participants and researchers) and were informed about how to consume them. The same testing procedure was repeated during the second session day (which took place at the same time as the first session), after the 28-day intervention period (average number of days between first test day and start of supplementation (SD): 8.5 (5.5); average number of days between end of supplementation and second test day (SD): 1.5 (0.7)). The second testing day ended with the written question to the participants on whether they thought they had taken the placebo or the probiotic, on which they answered by chance (number of correct answers: 15/29 (51,7%) for the placebo group and 12/29 (41.4%) for the probiotics group).

**Figure 1.**
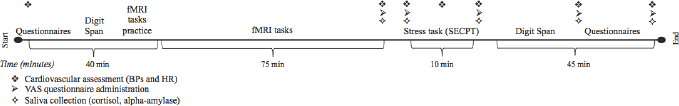
Overview of the testing session. Each participant was tested twice, before and after 4 weeks of supplementation with probiotics/placebo. The procedure of the two sessions was the same (i.e. participants performed the same tests in the same order). SECPT: socially evaluated cold pressor test; BP: blood pressure; HR: heart rate; VAS: visual analog scale.

#### 2.3.2 Questionnaires

Questionnaires were administered using an Electronic Data Capture (EDC) application for online data collection (Castor EDC, https://castoredc.com). Specifically, we assessed depression with the Dutch version of the self-reported Beck Depression Inventory questionnaire (BDI) (Beck, 1976). The 21 items of the BDI are rated on a 4-point Likert scale (from 0 to 3 per item) indicating the severity of the feeling of the participant in the past two weeks. A total BDI score between 0-13 is an indicator of no-minimal depression. A BDI score above 13 (i.e. 14-19: mild depression; 20-28: moderate depression; 29-63: severe depression) was an exclusion criterion. Depression sensitivity was evaluated with the Leiden Index of Depression Sensitivity-revised questionnaire (LEIDS-r) (Antypa et al., 2010), on which the effects of this specific probiotic product have already been reported previously (Steenbergen et al. 2015). Before completing the LEIDS-r, participants were asked to imagine a situation when they felt sad and to indicate, on a 5-point Likert scale ranging from 0 (i.e. ‘not at all’) to 4 (‘very strongly’), the degree to which each statement of the 34 items applied to them. The LEIDS-r total score was then calculated by adding the scores of its six sub-scales: aggression, control, hopelessness, risk aversion, rumination and acceptance. Other questionnaires were assessed to control for baseline differences in psychological traits, but these were not observed and, hence, reported in the supplement (Supplementary Materials).

#### 2.3.3 Task paradigms

The fMRI tasks (Figure 2) included an emotional face-matching paradigm, an emotional face-word Stroop paradigm, and the classic color-word Stroop paradigm. Participants were instructed to react as fast and accurately as possible during all three tasks. The experiments were programmed in Presentation^®^ software (Version 0.70, www.neurobs.com).

##### 2.3.3.1 Emotional face-matching paradigm

This paradigm (Figure 2*a*) was chosen to investigate intervention-induced changes in emotion reactivity (Hariri et al., 2000). Stimuli were presented in a block design, with a total of 18 blocks consisting of three stimuli each. The task included a control and an emotion condition. In the control condition participants had to match one of two geometric shapes presented at the bottom, to a target shape presented at the top of the screen. The experimental condition involved participants choosing one of two emotional (angry or fearful) faces presented at the bottom of the screen that best matched the emotional expression of a face seen at the top of the screen. The condition was kept constant over the block duration of 17 seconds, but was randomized between blocks. The total duration of the task amounted to seven minutes.

##### 2.3.3.2 Emotional face-word Stroop paradigm

A Dutch version of the emotional face-word Stroop task (Etkin et al., 2006) was used to assess intervention-induced differences in the ‘resolution of emotional conflicts’ in the face of emotional distracters (Figure 2*b*). During this task, participants were presented with pictures of male faces expressing fear or happiness. On top of the faces, the Dutch words for happy (i.e. “blij”) and fearful (i.e. “bang”) were presented in prominent red letters. The emotions described by the words were either congruent with the emotion of the face or incongruent, and participants had to indicate the emotion of the face by ignoring the emotion word. A total of 148 stimuli of happy or fearful faces were presented. The order of stimulus presentation was pseudo-randomized and the total duration of the task added up to 15 minutes.

##### 2.3.3.3. Classic color-word Stroop Paradigm

A Dutch version of the classic color-word Stroop task (Stroop, 1953) was used to assess intervention-induced differences in general cognitive control (i.e. resolving response conflict) in the absence of emotional stimuli (Figure 2*c*). During this task, participants were presented with four different color words written either in the same ink color as the word (e.g. red written in red ink) or in an incongruent color (e.g. red written in blue ink). They were asked to indicate the ink color of the word by pressing a button mapped to that color, and to ignore the word meaning. The task consisted of 80 stimulus presentations in total. Color-button mappings were randomized across subjects, but kept constant between the two sessions of each participant (for both Stroop tasks). The duration of the task amounted to 10 minutes.

**Figure 2.**
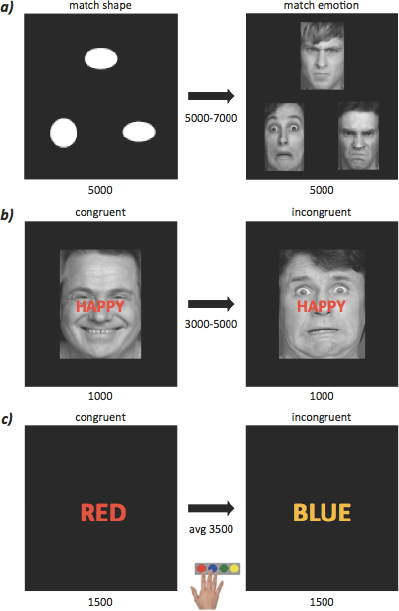
The fMRI paradigms. ***a)*** Emotional face-matching task, ***b*)** Emotional face-word Stroop task, and ***c)*** Color-word Stroop task. The control and the experimental conditions of each paradigm are represented on the left and right side of the panel respectively. The duration of each stimulus and the relative inter-trial interval are indicated in ms.

#### 2.3.4 Stress-induced working memory performance

For stress induction, we used the Socially Evaluated Cold Pressure Test (SECPT; Lovallo, 1975). During this test, physical and psychological stress was induced. Physical stress was induced by having participants immerse their hand into ice water (ranging between 0 and 3 degrees Celsius) for “as long as possible, until the researcher indicates to pull the hand out of the water”. Unknown to the participants, the maximal duration was set at three minutes (180 seconds). The mean duration of ice water immersion was 160.8 seconds (SD: 45.6) for the first session, and 171 seconds (SD: 32.4) for the second one. The test was conducted by a researcher who was yet unknown to the participant, and who adopted neutral and socially distant behavior to increase psychological stress. Further psychological stress was induced by asking participants to look into a video camera during the cold-water test, with the aim to record their facial expressions. Saliva samples were collected to evaluate cortisol and alpha-amylase levels in response to the stressor. A total of five saliva samples from each participant was obtained: one sample was obtained 10 minutes before the start of the SECPT, one sample right before and one sample immediately after the end of the SECPT, one sample 25 minutes and one sample 45 minutes after the end of the ice water immersion (see Figure 1). Parameters reflecting autonomic nervous system activation, i.e. systolic and diastolic blood pressure and heart rate (HR) were registered at the beginning of the test, as well as every time that the saliva samples were collected (plus one extra measurement between the collection of the second and third saliva sample). The total score of the visual analog scales (VAS) was used to assess the subjective feeling of stress and obtained by the sum of each sub-scale of the VAS: tension, happiness (reversed scored), pain, fear, irritation, and stress. The VAS questionnaires were completed each time during saliva collection.

To evaluate stress-related effects in cognitive functioning, we used the digit span test to assess working memory performance, at the beginning of the experiment and right after the stress induction (within a range of 5 minutes from the end of the stress task). Specifically, during the digit span test participants listened to a series of numbers and tried to repeat each series correctly (DS forward) or repeat it backwards (DS backward). Following a correct response, the participants had to repeat increasingly longer sequences. Participants got different versions of the test before and after stress on the first and second test day.

### 2.4 Data analyses

#### 2.4.1 Questionnaire analysis

Statistical analyses of the data were performed using IBM SPSS statistics (version 23.0), and the results were expressed in terms of mean values and standard errors of the mean (SEM). The effects of the intervention on questionnaire scores were analyzed by performing 2×2 repeated-measure ANOVAs for data that was normally distributed. The first factor was ‘Group’ (between-subjects), with two levels (placebo and probiotics), the second factor was ‘Session’ (within-subjects), with two levels (pre- and post-intervention session). For data that was not normally distributed, we used the Wilcoxon-Mann-Whitney test to assess group differences on the post-pre scores.

#### 2.4.2 Behavioral analysis of fMRI tasks

We evaluated behavioral performance during the fMRI tasks before and after the intervention between the two groups, by analyzing the correct reaction times (RTs). The analyses were done on log-transformed data. We ran a 2×2×2 repeated-measure ANOVA design, with the factors Group (between-subjects), Session (within-subjects), and Condition (experimental vs. control condition, within-subjects).

#### 2.4.3 fMRI acquisition and analyses

##### 2.4.3.1 MR data acquisition

MR data were acquired using a 3T MAGNETOM Prisma system, equipped with a 32-channel head coil. During the three tasks, 3D echo planar imaging (EPI) scans using a T2*weighted gradient echo multi-echo sequence (Poser et al., 2006) were acquired (voxel size 3.5 × 3.5 × 3 mm isotropic, TR = 2070 ms, TE = 9 ms; 19.25 ms; 29.5 ms; 39.75 ms, FoV = 224mm). The slab positioning and rotation (average angle of 14 degrees to AC axis) optimally covered both prefrontal and deep brain regions (i.e. including affective brain regions like the amygdala). A whole-brain high-resolution T1-weighted anatomical scan was acquired using a MPRAGE sequence (voxel size 1.0 × 1.0 × 1.0 isotropic, TR = 2300 ms, TE = 3.03 ms, 192 slices).

##### 2.4.3.2 fMRI data preprocessing

Data was preprocessed and analyzed using Statistical Parametric Mapping (SPM8) (Wellcome Department of Imaging Neuroscience, London). Volumes for each echo-time were realigned using six rigid body spatial transformations (translations and rotations: x, y, z, pitch, roll, jaw). Thirty volumes acquired before the tasks were used to combine the four echo images into a single volume using an echo weighting method known as PAID-weighting (Poser et al., 2006). Resulting combined functional (EPI) images were slice-time corrected by realigning the time series for each voxel to the time of acquisition of the reference slice. Subject-specific structural and functional data were subsequently co-registered to a standard structural or functional stereotactic space respectively, using Montreal Neurological Institute (MNI) templates. A unified segmentation approach was then used to segment the structural images. Segmented images were subsequently spatially co-registered to the mean of the functional images. The transformation matrix resulting from the segmentation step was used to normalize the structural and functional images to MNI space, resampled at a voxel size of 2 × 2 × 2 mm. In a final step, normalized functional images were spatially smoothed using an 8 mm full-width at half maximum (FWHM) Gaussian kernel.

##### 2.4.3.3 fMRI analyses

Fixed effects analyses of the emotional face-matching paradigm were carried out at the first level using a block-design fMRI approach (i.e. 12 ‘emotion’ blocks and 6 ‘shape’ blocks of each 17 seconds). Two regressors of interest were compared: ‘emotion’ minus ‘shape’ condition. Similar first level analyses were performed using an event-related approach for the Stroop paradigms. For both the emotional face-word Stroop paradigm and the color-word Stroop paradigm the statistical model contained two regressors of interest, which we subtracted for our contrast of interest: ‘incongruent’ minus ‘congruent’ condition. Missed and incorrect trials were taken into account in a regressor of non-interest for both paradigms. Additionally, thirteen regressors of non-interest were added to the designs of all three tasks, including twelve rigid-body transformation parameters (i.e. movement regressors consisting of three translations, three rotations and their linear derivatives) obtained during realignment, as well as one constant term. A high-pass filter with a cut-off of 128 seconds was applied to the time-series of the functional images to remove low-frequency drifts. By applying an autoregressive AR (1) model, correction for serial correlations was carried out. Both sessions of each subject were included in one first level model.

On the second (group) level, we first performed random effect analyses of variance (ANOVA) in a full-factorial design to obtain the main task effects (positive and negative) for each task across sessions. The ANOVA analyses were run with the contrast images specified in the first level analyses and two additional factors: Group (probiotics and placebo) as a between subject-factor and Session (pre- and post-intervention session) as a within-subject factor. Subsequently, we ran two-samples t-test analyses between the probiotics and the placebo group using the contrast images of Condition x Session specified at the first level. We considered results significant if p<.05 Family-Wise-Error (FWE) whole-brain corrected at cluster level (with a cluster defining threshold of p<.001).

#### 2.5.3 Stress-related data analyses

Using a 2×2×2 repeated-measure ANOVA, we analyzed the stress-related cardiovascular data, i.e. systolic (BPsys) and diastolic blood pressure (BPdia), as well as heart rate (HR); hormones, i.e. alpha-amylase and cortisol; and VAS scores (subjective stress-related feeling). All the analyses were done on log-transformed data. The first factor was Group, with two levels (placebo and probiotics, between-subjects), the second factor was Session, with two levels (pre- and post-intervention session, within-subjects), and the third factor was Time (within-subjects). For the cardiovascular data, the third factor Time consisted of seven levels (one level for each saliva collection time point (see above) plus two extra measurements: one at the beginning of the experiment, and one right before the immersion of the participant’s hand in the cold water box), while for the alpha-amylase, cortisol, and VAS scores the factor Time consisted of five levels (one level for each saliva collection time point; see Figure 1). Effects of Time indicated an effect of the stressor, whereas Group*Session interactions indicated effects of probiotic supplementation.

A 2×2×2 repeated measure ANOVA was also chosen to analyze probiotics-induced effects (Group*Session interactions) on DS backwards after versus before stress induction (within-subjects factor: Time). Analyses were performed on the raw scores given that the DS scores were normally distributed.

In the last step of the analysis, we investigated whether individual differences in probiotics effects on working memory were associated with probiotics effects on brain functioning during cognitive control (i.e. incongruent versus congruent Stroop trials). For this, we extracted averaged beta weights (MarsBar toolbox of SPM, Brett, 2002) from the significantly (pFWE<.05) activated regions in frontal cortex during the Stroop task (incongruent>congruent, Figure 3f), independent of the intervention. To extract the average beta values, we applied a sphere of 10 mm on the local maxima of each (sub-)cluster in frontal cortex, to prevent averaging from large clusters extending across different regions. Subsequently, we calculated post- minus pre-intervention scores of the average beta weights and correlated these to the intervention effect (post- minus pre-intervention) in the stress-induced working memory (i.e. DS backward) performance scores. We performed these ROI analyses per group (probiotics and placebo), such that we could compare the correlations of the probiotics group to that of the placebo group using Fisher’s r to z transformation.

We considered results with p<.05 significant or, with multiple comparisons with the same outcome (i.e. correlation of intervention effects on stress-related working memory in multiple frontal Stroop ROIs), p<.05/number of comparisons (Bonferroni correction). We report partial eta squared as a measure of effect size.

## 3. Results

### 3.1 Questionnaires

Table 1 shows the scores on the questionnaires before and after the intervention for the placebo and probiotics group. To assess the effects of probiotics on depression and depression sensitivity, we assessed the interaction between the factors Group and Session. We did not observe effects of probiotics on any of the questionnaires (Supplementary Materials). We also did not find differences between the probiotics and placebo group at baseline, i.e. pre-intervention (all p>.05).

**Table 1:** Raw scores on the questionnaires expressed as mean values (SEM).

|  | PLACEBO | PLACEBO | PROBIOTICS | PROBIOTICS | Group*Session |
| --- | --- | --- | --- | --- | --- |
|  | pre | post | pre | post | p-values |
| *BDI* | 2.62 (0.54) | 2.24 (0.58) | 2.10 (0.37) | 1.78 (0.39) | .95 |
| *LEIDS-r* | 28.59 (2.58) | 21.66 (2.08) | 29.97 (2.35) | 27.24 (2.55) | .09 |

### 3.2 RT results

First, we assessed the main effect of the task conditions across sessions and groups in terms of RTs (see raw data in Table 2 and Figure 3 a), c), and e)). As expected, in the emotional face-matching task, participants were significantly slower in the ‘emotion’ than in the ‘shape’ condition (main Condition: F(1,56) = 654.65, p< .001, η^p2^=.921). Similarly, during the two Stroop tasks, participants were significantly slower in the incongruent than in the congruent conditions (emotional face-word Stroop, main Condition: F(1,56) = 276.60, p<.001, η^p2^=.921; color-word Stroop, main Condition: F(1,56) = 231.05, p<.001, η^p2^=.805). Subsequently, we assessed the effects of the probiotics versus placebo on the three tasks, but we did not observe significant Group x Session x Condition effects (all p>.05, all η^p2^<.005). We also did not find differences between the probiotics and placebo group at baseline, i.e. pre-intervention (all p>.05).

**Table 2:** Mean (SEM) RTs for each fMRI paradigm, before and after the intervention with probiotics and placebo.

| Paradigm | Condition | PLACEBO | PLACEBO | PROBIOTICS | PROBIOTICS |
| --- | --- | --- | --- | --- | --- |
|  |  | pre | post | pre | post |
| Emotional<br>face-matching | Shape | 867.85<br>(36.9) | 775.18<br>(27.29) | 821.46<br>(22.50) | 779.03<br>(25.89) |
|  | Emotion | 1352.73<br>(57.67) | 1282.57<br>(54.56) | 1364.69<br>(68.81) | 1253.71<br>(59.53) |
| Emotional<br>face-word<br>Stroop | Congruent | 778.44<br>(29.85) | 795.55<br>(37.72) | 761.60<br>(41.50) | 762.11<br>(33.39) |
|  | Incongruent | 829.63<br>(30.63) | 847.29<br>(39.55) | 825.89<br>(44.18) | 812.06<br>(33.01) |
| Color-word<br>Stroop | Congruent | 856.71<br>(42.21) | 817.61<br>(35.09) | 801.45<br>(52.49) | 784.00<br>(38.22) |
|  | Incongruent | 974.80<br>(44.67) | 955.49<br>(43.02) | 917.32<br>(52.07) | 893.74<br>(51.26) |

*No significant Group x Session x Condition effects*

### 3.3 Neuroimaging results

The main task effects of the three fMRI tasks are shown in Figure 3 b), d), and f), and Table 3 at pFWE<.05 (whole-brain correction at cluster level). The emotional face-matching task activated, amongst others, the bilateral amygdala. We observed responses to the emotional face-word Stroop task in regions such as the medial frontal cortex (pre-SMA), vmPFC and lateral PFC. Clusters activated for the color-word Stroop task included the lateral PFC and medial frontal cortex.

**Figure 3.**
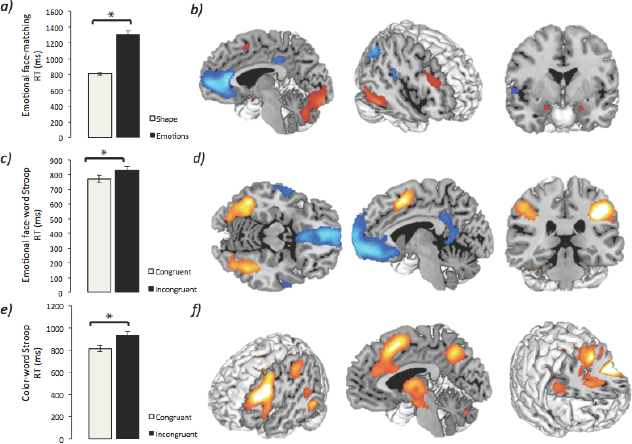
Main tasks effects. Main tasks effects (i.e. across groups and sessions) for **a)** the emotional face-matching paradigm in RTs and **b)** in brain (hot colors: emotion > shape; cold colors: shape > emotion). Main tasks effects for **c)** the emotional face-word Stroop paradigm in RTs and **d)** in brain (hot colors: incongruent > congruent; cold colors: congruent > incongruent). Main tasks effects for **e)** the color-word Stroop paradigm in RTs and **f)** in brain (hot colors: incongruent > congruent). Clusters are significant at pFWE<.05, cluster defining threshold: p<.001.

**Table 3:**
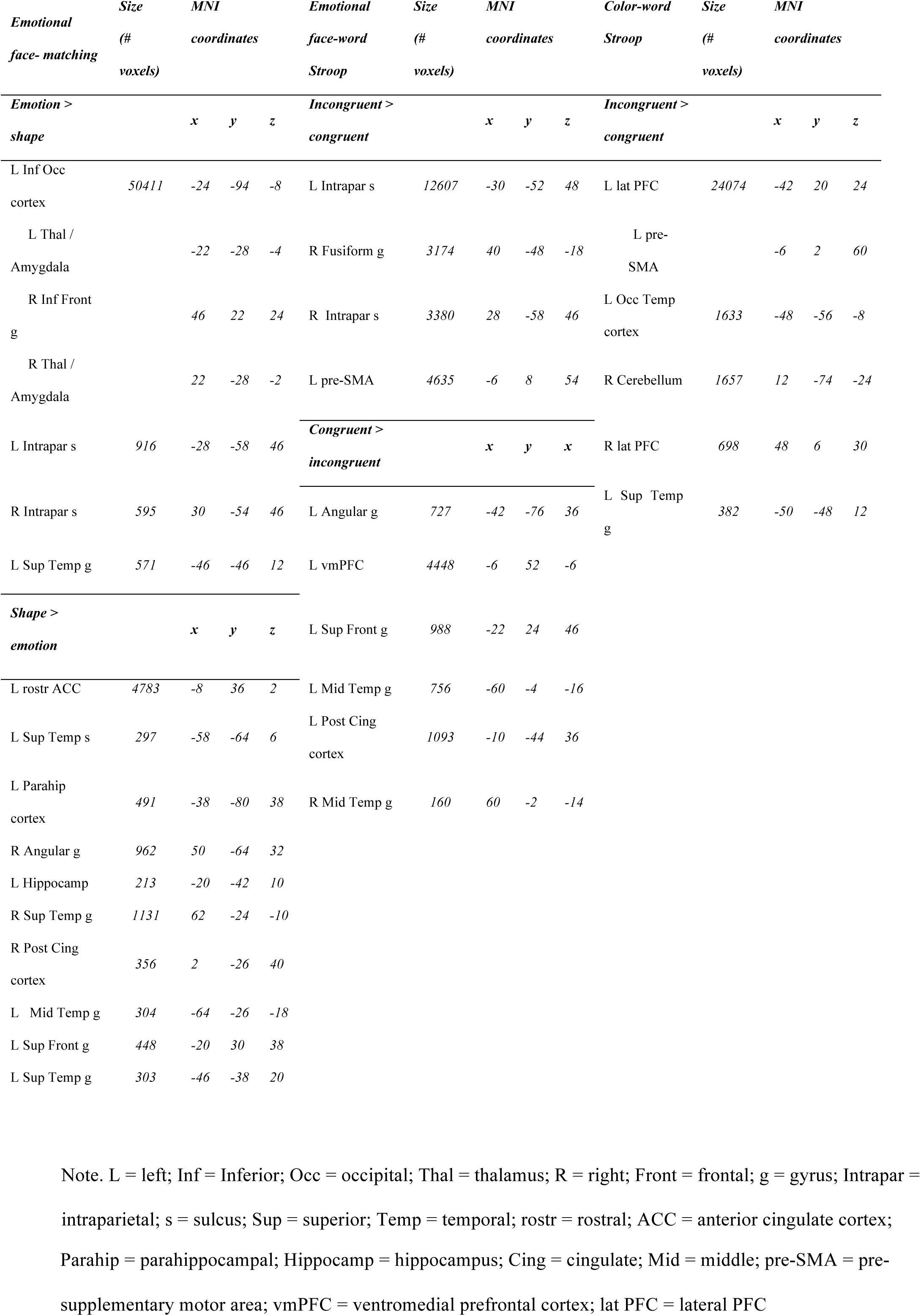
*Main task activations at P_FWE_<.05 (cluster level) across sessions and groups.*

We subsequently assessed the effects of probiotics versus placebo (pre vs. post intervention) on each of the three fMRI tasks. At our whole-brain corrected threshold of p_FWE_<.05 (cluster level), we did not observe any effects of probiotics during emotional face-matching, emotional face-word Stroop or color-word Stroop.

### 3.4 Stress-related results

#### 3.4.1 Stress-induced working memory performance

The second aim of the present study was to investigate the effects of probiotics on stress-related changes in working memory performance. First, we assessed whether the SECPT indeed induced stress by measuring cardiovascular parameters (BPsys, BPdia, and HR), hormones (cortisol: probiotics n=26 and placebo n=27; alpha-amylase: probiotics n=21 and placebo n=20), and subjective stress-related feelings (VAS total score: probiotics n=28 and placebo n=29).

Significant effects of Time revealed that the stressor indeed had effects on physiological and subjective measures of stress (all p < .05) (Supplementary Materials) see Figure 4a and 4b. However, the intervention did not affect these physiological and subjective measures of stress (Group x Session x Time interactions for HR, BPsys, BPdia, hormones, and VAS: p > .05) (Supplementary Materials).

**Figure 4.**
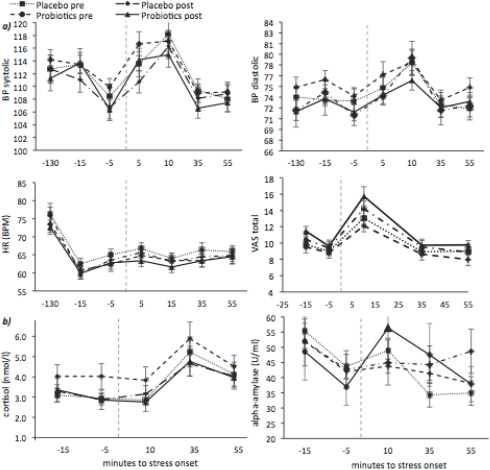
Physiological and psychological stress effects. ***a)*** Physiological and psychological raw data during the SECPT (onset at time = 0, dotted vertical line), i.e. blood pressure (BP) systolic (top-left panel), blood pressure (BP) diastolic (top-right panel), heart rate (HR) values (bottom-left panel), and total VAS score (bottom-right panel) and **b)** Hormone levels during the SECPT, i.e. cortisol values (left panel), and alpha-amylase values (right panel), for the two groups (probiotics and placebo), before and after the intervention period.

We were specifically interested in the effects of the stressor on working memory (DS backward) performance and its potential modulation by probiotics. Raw digit span scores and SEMs are reported in the Supplementary Materials.

We found that stress-induced working memory performance in DS backward was differentially affected by the probiotics (post vs. pre) than by placebo (post vs. pre), evidenced by a Time(2) x Group(2) x Session(2) interaction (F(1,56)=4.48, p=.039, η^p2^=.07). Breaking this interaction effect down into simple effects, we observed that the probiotics (post vs. pre) tended to increase stress-induced backward digit span performance (Time(2) x Session(2), probiotics: F(1,28)=4.1, p=.053, η^p2^=.127), whereas no significant post-pre intervention effect was observed in the placebo group (Time(2) x Session(2), placebo: F(1,28)<1, p=.36, η^p2^=.03) (Figure 5). We did not find differences between the probiotics and placebo group at baseline, i.e. pre-intervention differences in the effect of stress on DS backwards (Time(2) x Group(2), pre-intervention: F(1,56)=3.24, p=.077, η^p2^=.05). In sum, probiotics versus placebo supplementation resulted in a buffer against the detrimental effect of stress on control-demanding working memory.

**Figure 5.**
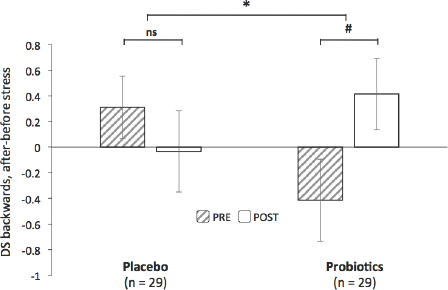
Stress-induced changes in working memory. Stress-induced working memory scores (calculated as the difference of DS backwards scores after stress minus scores before stress) obtained pre- and post-intervention for the placebo and for the probiotics group. * = p<.05; # = p<.1; ns = p>.1

#### 3.4.2 Individual differences in the effects of probiotics: Correlations between neural cognitive control responses and stress-induced working memory effects

We extracted averaged beta weights from the three frontal cortex clusters that showed increased responses for the main effect of the color-word Stroop task (incongruent > congruent), i.e. independent of intervention effects to avoid circularity (see Figure 3f). Subsequently, we calculated the post- minus pre-intervention difference score of these averaged beta weights per independent ROI and correlated them to the intervention effects in the stress-related working memory scores per group. In the probiotics group, we found significant negative correlations between the intervention effect on the averaged incongruent-congruent betas and the intervention effect on the stress-related difference scores in DS backwards in all three frontal ROIs (see Table 3 for the clusters) activated during the color-word Stroop task (left lateral PFC (BA45) (x,y,z: −42, 20, 24): r = −0.52, p=.004; pre-SMA (x,y,z: −6, 2, 60): r = −0.38, p=.04; right lateral PFC (BA44) (x,y,z: 48, 6, 30): r = −0.60, p<.001) (Figure 6). However, in contrast to the two lateral PFC regions, the association in pre-SMA was not significant after correction for multiple comparisons (p> 0.017). See Supplementary Materials for the fronto-striatal responses observed in the whole-brain correlation within the probiotics group. These brain-behavior correlations were not found in the placebo group (left lateral PFC (BA45): r = −.21, p=.27; pre-SMA: r = −0.19, p=.32; right lateral PFC (BA44): r= 0.03, p=.89) (Figure 6). Importantly, we assessed whether the brain-behavior correlations were significantly different between the probiotics and placebo group. Indeed, the intervention with probiotics resulted in a greater association between changes in stress-related working memory and neural cognitive control responses in the right lateral PFC than the placebo intervention (Fisher’s *r* to *z* transformation, z= −2.61, p= .009). In contrast to the right lateral PFC, the correlation coefficients between the probiotics and placebo groups did not significantly differ from each other in the left lateral PFC (z=-1.31, p=.19) or in the pre-SMA (z=-0.75, p=.45).

**Figure 6.**
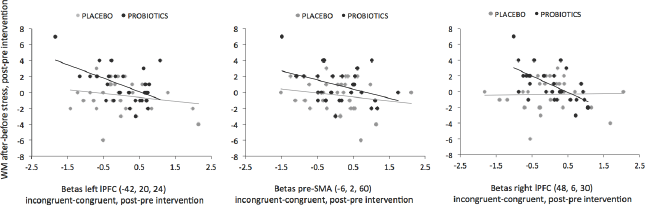
Brain-behavior correlations. Correlations between the intervention effect on stress-induced working memory (WM) difference scores (digt span backwards scores after stress minus digit span backwards scores before stress, post-pre intervention) and the intervention effect on cognitive control-related brain responses (i.e. averaged beta values for incongruent-congruent color-word Stroop trials, post-pre intervention) for each independently selected frontal ROI (see Figure 3f and Table 3), i.e. from left to right: left lateral PFC (BA45), pre-SMA, and right lateral PFC (BA44) for the placebo (grey dots) and for the probiotics group (black dots).

In sum, the probiotics’ intervention effect (post-pre) on stress buffer during working memory was especially evident in those subjects with probiotics’ induced decreases in prefrontal cortex recruitment during cognitive control. This brain-behavior association in right lateral PFC was significantly greater for the probiotics than for the placebo group.

## 4. Discussion

We aimed to investigate the neurocognitive mechanisms of multi-species probiotic supplementation in healthy human volunteers. Specifically, for our first aim, we assessed whether the neurocognitive effects of probiotics versus placebo would only be seen on emotion reactivity to negative stimuli or also on emotion regulation or general cognitive control. In our secondary aim, we were interested in the effects of probiotics on stress-induced changes in working memory and associated neural control mechanisms. Four weeks of probiotic supplementation did not result in differences relative to placebo in terms of neural or behavioral responses during emotion reactivity, emotion regulation or cognitive control. However, the groups did differ in their cognitive response to an acute stressor; i.e., the probiotics group showed an increased buffer against the negative effects of stress on working memory performance relative to placebo. This increased cognitive buffer against stress was especially seen in individuals with probiotic-induced decreases in prefrontal cortex recruitment during cognitive control and this association differed significantly from the one in the placebo group.

All three fMRI paradigms robustly activated the expected brain regions and showed the expected behavioral effect. Specifically, emotion reactivity was seen for instance in the amygdala and in terms of RTs during emotional face matching (Hariri et al., 2000; Haxby et al., 2000), while emotion regulation and general cognitive control was observed in frontal regions and in longer RTs during the two Stroop tasks (Etkin et al., 2006; Roberts and Hall, 2008)(Aarts et al., 2008, 2009; Cieslik et al., 2015; Roberts and Hall, 2008). However, we did not find significant effects of probiotic supplementation on the behavioral (i.e. RTs) and neural responses to the tasks. These neurocognitive results are in line with the results on the questionnaires, which also revealed no differences between probiotics and placebo. This contrasts with previous findings in healthy controls, using the same probiotic product, which indicated a probiotics-induced decrease in cognitive reactivity to sad mood using the LEIDS-r questionnaire (Steenbergen et al., 2015). However, when comparing the baseline scores on the depression-related questionnaires, i.e. LEIDS-r and BDI, we note that our participants scored much lower (i.e. across groups, mean LEIDS-r: 25.1; mean BDI: 2.4) than the participants of Steenbergen and colleagues (i.e. across groups, mean LEIDS-r: 43.7; mean BDI: 8.5). Recently, a study in IBS patients also found a probiotics-induced reduction of depressive symptoms as well as a reduction in neural emotion reactivity using fMRI, but these patients had mild to moderate depression at baseline (i.e. across groups, mean HADS-depression: 10.5) (Pinto-Sanchez et al., 2017). Accordingly, studies assessing probiotics’ effects in participants with at least some degree of depression at baseline (i.e. across groups, median HADS-depression: 5.5) (Messaoudi et al., 2011) or in patients with major depressive disorder (Akkasheh et al., 2016) have reported positive effects on depression scales. In line with our results, the sample of Tillisch and colleagues (2013) scored low on depression (i.e. across groups, mean HADS-depression: 1.3) and they did not observe differences between the placebo and probiotics group on self-report or neural measures of emotion reactivity (neural differences were only observed for an ill-controlled comparison between probiotics and no intervention). Similarly, another study with a healthy sample with low depression ratings (mean BDI at baseline: 3.92) did not find probiotics-induced changes in cognition or resting EEG (Kelly et al., 2017). Healthy individuals are known to exhibit a different gut-microbiome composition relative to individuals suffering from depression (Jiang et al., 2015; Kelly et al., 2016; Naseribafrouei et al., 2014). Thus, based on our current and on previous results, it seems that probiotics – compared with placebo - only have effects on self-report and neural measures of emotion and cognition if subjects are either clinically affected or score high in diagnostic questionnaires, suggesting limited beneficial effects of probiotics on mood and neurocognition in healthy individuals (i.e., free of psychological, endocrine, and gastro-enteric diseases and with no drug-, severe alcohol-, or prolonged medication use) as in the current study.

However, one study, using *Bifidobacterium longum* 1714, did demonstrate probiotic-induced beneficial effects on associate learning and changes in resting EEG in a healthy, low depressed sample (i.e. mean BDI at baseline: 3.6) (Allen et al., 2016). The same strain was able to reduce anxiety and depression-like behavior in anxious BALB/c mice (Savignac et al., 2014). Perhaps, in addition to the presence of a certain degree of depressive/anxiety symptoms, other factors such as type of bacterial strains play a role in observing beneficial neurocognitive effects of probiotics. Therefore, future studies should investigate the mechanisms of action of single bacterial strains to understand their potential in benefitting mood and cognition in different human populations.

The second aim of the present study was to investigate whether probiotics can have a beneficial effect on cognition by buffering against stress. Our results demonstrated an increase in working memory performance after stress induction in those participants who consumed probiotics, which was also reflected in associated probiotics-induced neural changes during cognitive control. The cardiovascular, hormonal, and psychological measures of stress indicated that stress was reliably induced, but were themselves not influenced by the probiotics, which is in line with previous studies (Kelly et al., 2017; Mohammadi et al., 2016; Moller et al., 2017) but see (Allen et al., 2016). The finding that probiotic supplementation did affect cognitive control, i.e. working memory performance, as a function of stress, is in line with preclinical work emphasizing the role of gut microbiota and probiotics in stress-related disorders (for a review, see e. g. Kelly et al., 2015). For example, in mice with a bacterial infection, acute stress caused memory impairments, which were ameliorated by probiotics (Gareau et al., 2011). Moreover, *Bifidobacterium longum* was able to improve learning and memory in mice with high susceptibility to stress (Savignac et al., 2015). Based on our results, we hypothesize that in healthy young individuals, probiotics might especially induce beneficial effects on cognition and brain functioning when the system is challenged by stress.

The buffer against stress-induced detriments in working memory in the probiotics group was especially seen in individuals with probiotics-induced changes in frontal brain regions during incongruent versus congruent trials in the Stroop task. Individual working memory capacity is predictive of Stroop performance (Kane and Engle, 2003). Thus, it might not be surprising that we observed correlations between the probiotics’ effects in both cognitive control measures. Both working memory and Stroop conflict are known to elicit neural responses in the frontal cortex, such as in the lateral PFC, which is important for goal-directed behavior (Aarts et al., 2008; Brass and von Cramon, 2004; Garavan et al., 2002; Miller and Cohen, 2001). Increases in probiotics-induced protection against stress effects on working memory were related to probiotics-induced decreases in frontal cortex recruitment during cognitive control. In cognitive control tasks, such as the color-word Stroop task, the frontal cortex is generally more recruited with increasing difficulty. Here, diminished frontal cortex responses were observed in the absence of behavioral changes, which has previously been interpreted as more efficient frontal cortex functioning (see also e.g. Mattay et al., 2003). Importantly, we did not observe significant correlations in the placebo group and the effects in the right lateral prefrontal cortex were significantly greater in the probiotics than in the placebo group. This means that the brain-behavior association with stress–related working memory was specific to the probiotics treatment and could not be present due to general factors that would change post- versus pre-intervention for both groups, such as those related to practice for instance. The placebo group as a whole seemed to demonstrate (*non*-significantly) reduced detrimental effects of stress on working memory relative to the probiotics group at baseline, which could have resulted in ceiling effects, i.e. not enough room for improvement. However, the lack of brain-behavior correlations between stress-induced working memory and cognitive control brain responses in the placebo group cannot easily be explained by ceiling effects given the focus on individual differences.

Nevertheless, the biological mechanisms behind these effects remain to be elucidated, particularly as the beneficial effects of probiotics on stress-related cognition were not accompanied by probiotics-induced changes in HPA-axis (i.e. cortisol) or sympatho-adreno-medullary system (i.e. alpha-amylase) markers. If the current multi-species probiotic product indeed strengthened epithelial barrier function, as demonstrated *in vitro* (Van Hemert, 2014), then other mechanisms could underlie the currently observed effects. Stress can increase the permeability of the intestinal barrier and subsequent immune reactions to LPS crossing the barrier. Indeed, acute stress paradigms can increase levels of the cytokine interleukin-6 in healthy human volunteers (Treadway et al., 2017). Moreover, multi-species probiotics have been shown to reduce inflammatory markers in depressive patients (Akkasheh et al., 2016) and in patients with type 2 diabetes (Asemi et al., 2013). In reaction to increased levels of LPS, pro-inflammatory cytokines can enter the central nervous system and negatively influence brain processes involved in memory and learning (Capuron and Miller, 2011; Miller, 1998; Rogers et al., 2016; Sparkman et al., 2006). Working memory is particularly modulated by signaling of the neurotransmitter dopamine in frontal and striatal brain regions (Cools and D’Esposito, 2011), and dopamine neurotransmission is affected by stress (Arnsten and Goldman-Rakic, 1998; Bliss et al., 1968). Hence, acute stress is generally known to be detrimental to working memory performance and working memory-related brain responses in PFC, particularly in tasks requiring modulation of information in working memory, such as the digit span backward and the n-back task (Qin et al., 2009; Schoofs et al., 2009). Increases in inflammatory tone in the body can induce neuro-inflammation, which can particularly affect dopamine signaling, as shown in humans and in non-human primates (Felger and Treadway, 2017). Probiotics might increase the buffer against stress-induced reductions in dopamine-dependent working memory performance by decreasing (stress-induced) permeability of the intestinal barrier, reducing blood concentration of LPS, and reducing brain levels of pro-inflammatory cytokines, as shown previously in rats (Ait-Belgnaoui et al., 2012).

An alternative mechanism of the probiotics-induced beneficial effects might be through the production of metabolites. For example, the gut microbiome has the potential to synthesize precursors of the monoamine neurotransmitters (i.e. large neutral amino acids) that could enter the blood stream, cross the blood-brain barrier, and affect neurotransmitter release (Lyte, 2013; Sampson and Mazmanian, 2015). We have recently demonstrated that predicted microbial potential to synthesize phenylalanine, a precursor of dopamine, was associated with neural responses during reward anticipation (Aarts et al., 2017), a function that is also typically modulated by dopamine (Knutson and Gibbs, 2007). The influence of the gut microbiome on central dopamine processing is also evident from germ-free mice, who exhibit increased turnover of dopamine in the brain (Diaz Heijtz et al., 2011). Future studies should investigate the potential mechanisms of probiotics in affecting central neurotransmitter release, e.g. by precursor production, but also by short chain fatty acid production, or by signaling on enteric nerve cells and the vagus nerve (DeCastro et al., 2005; Mally et al., 2004; Sampson and Mazmanian, 2015).

## 5. Conclusions

We showed that 4-weeks of supplementation with a multi-species probiotic positively affected cognition under challenging situations induced by acute stress, which was associated with changes in frontal brain regions during cognitive control. However, on neurocognitive tasks administered in relative neutral situations – even if they contained processing of mild negative emotional information – we did not observe effects of probiotics across the group. Our findings of stress-dependent beneficial effects of probiotics on cognition can be of clinical importance for stress-related psychiatric and gastro-intestinal disorders.

### Funding sources

The study was supported by the Dutch Ministry of Economic Affairs under the TKI Life Science and Health, project LSHM15034, and by Winclove Probiotics B.V., The Netherlands. EA and JW were supported by a Food, Cognition and Behaviour grant from the Netherlands Organization for Scientific Research (NWO, grant 057-14-001). EA was also supported by a Veni grant from the Netherlands Organization for Scientific Research (NWO, grant 016.135.023). KR was supported by a starting grant from the European Research Council (ERC_StG2012_313749) and by a VICI grant (#453-12-001) from the Netherlands Organization for Scientific Research (NWO). SvH is employed at Winclove Probiotics B.V. This does not alter this authors’ adherence to all publication policies on sharing data and materials. All other authors have no conflicts of interest to declare.

## Supplementary Materials Papalini et al

### Stress matters: effects of a multispecies probiotic on neurocognitive measures of emotion, cognitive control, and working memory

#### Additional questionnaires

We assessed the affective response (sensitivity) to reward and punishment using the Behavioral Inhibition Scale (BIS) / Behavioral Approach Scale (BAS), (Carver and White, 1994). See Table S1 below.

**Table S1.**
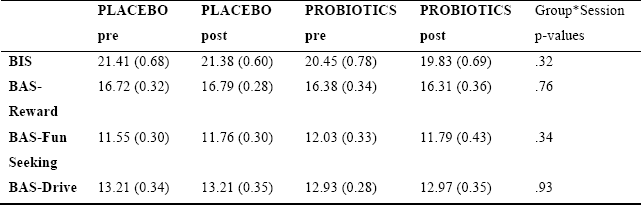
BIS-BAS questionnaire scores

#### Saliva and cardiovascular data

Participants were instructed not to consume food and beverages other than water or to exercise within the two hours preceding the start of the saliva collection. The saliva samples were collected via absorbent devices (salivettes -Sarstedt, Nümbrecht, Germany) and immediately frozen (at the temperature of −24°). Cortisol and alpha-amylase parameters were analyzed at the end of the experiment by an independent and specialized lab.

Cardiovascular monitoring, i.e. heart rate and blood pressure (systolic and diastolic), were assessed using a standard upper arm blood pressure monitor medical device.

#### Stress-related results

For the cardiovascular parameters BPsys and BPdia, we found significant effects of the stressor (main Time (7), BPsys: F(6,51) = 20.7, p <.001, η^p2^=.708; BPdia: F(6,51) = 8.3, p <.001, η^p2^=.493) and of repeating the stressor (Session(2), BPsys: F(1,56) = 6.01, p =.01, η^p2^=.097; BPdia: F(1,56) = 5.7, p = .02, η^p2^=.093). For both blood pressures we did not find a significant Time(7) x Group(2) x Session(2) interaction (BPsys: F(6,51)<1, η^p2^=.086 and BPdia: F(6,51)<1, η^p2^=.091), or any other significant interaction (all p>.05). For HR, cortisol, and alpha-amylase levels, we similarly observed an effect of the stressor (main Time (7) HR: F(6,51) = 21.9, p<.001, η^p2^=.721; main Time (5) cortisol: F(4,48) = 16.8, p <.001, η^p2^=.583; main Time (5) alpha-amylase: F(4,36) = 3.9, p =.01, η^p2^=.304), but no other main or interaction effects (all p>.05).

We calculated a total VAS score by adding up the six subscales (irritation, tension, happiness (reverse scoring), pain, fear, and stress levels) to obtain one measure of subjective feeling of stress. We found a significant effect of the stressor (Time(5): F(4,52) = 36.8, p <.001 η^p2^=.739) and of repeating it (Session(2): F(1,55) = 4.7, p = .035, η^p2^=.078), and a significant interaction between Time and Session (Time*Session: F(4,52) = 2.7, p = .04, η^p2^=.171); however, these effects did not differ across groups (all interactions with Group, p>.05). We also did not find differences between the probiotics and placebo group at baseline, i.e. pre-intervention, for any of the physiological or subjective stress variables (all p>.05) except for the sub-scale VAS ‘tension’ (p =.02) where the probiotics group indicated to feel more tense (SD): mean 2.79 (1.8), in comparison with the placebo group (SD): mean 1.93 (.99). No group differences at baseline were observed for the total VAS score. To sum up, although the stressor worked, the absence of interaction with Session and Group demonstrated that physiological and subjective stress measures were not affected by the probiotics.

#### Stress-induced working memory performance

As expected, the effects were specific to the cognitively more demanding digit span backward test (see main text), as we did not find a significant Time(2) x Group(2) x Session(2) interaction for digit span forward performance (F(1,56)<1). For raw digit span scores, see Table S2 below.

**Table S2.**
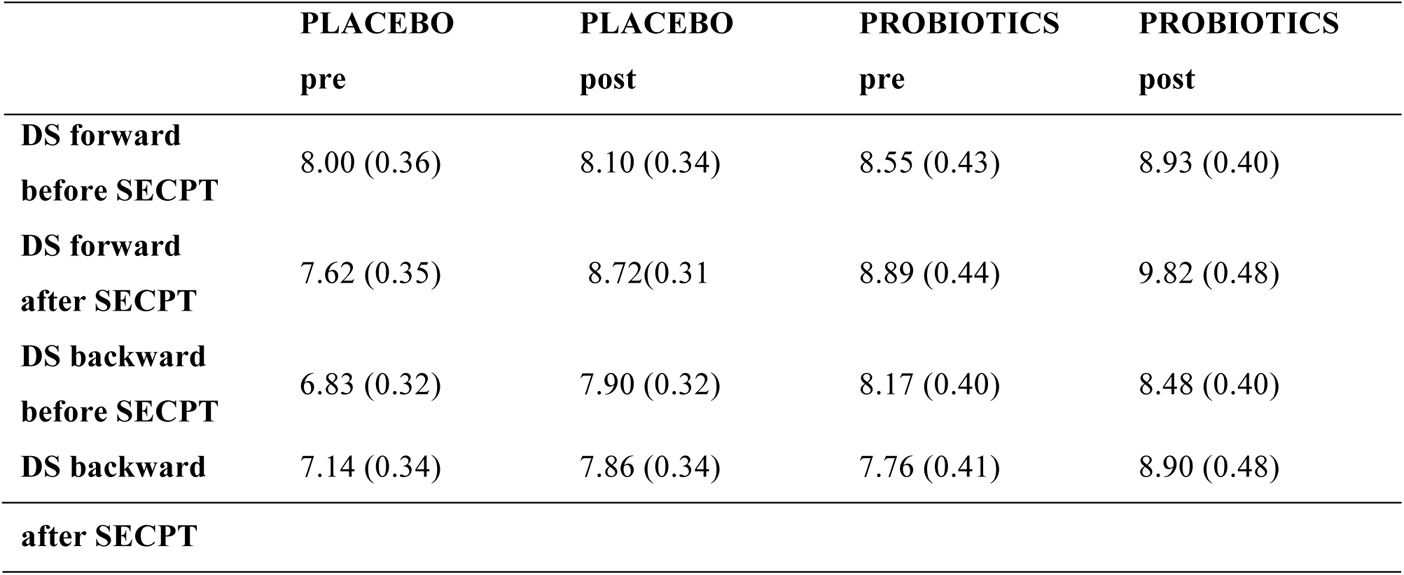
Mean (SEM) Digit Span scores (backward) before and after stress induction, and before and after the supplementation period with placebo or probiotics.

#### Correlations between neural Stroop responses and stress-induced working memory effects: whole-brain analysis

We used the post- minus pre-intervention stress-related working memory scores as a regressor of interest in the original t-test model of the fMRI data of the color-word Stroop task (i.e. incongruent > congruent, post- > pre-intervention), separately for each group. Only within the probiotics group, we found whole-brain corrected significant associations between probiotic-induced (post-pre intervention) increases in stress-related DS backwards and probiotic-induced decreases in brain responses (incongruent-congruent) during the color-word Stroop task in striatum, bilateral PFC, and medial frontal cortex (p_FWE_<.05 at cluster level, Table S3 + Figure S1).

**Table S3.** Whole-brain corrected significant clusters (p_FWE_<.05, cluster defining threshold: p<.001) activated during the color-word Stroop task (post-pre intervention) that were correlated with stress-related changes in working memory in the probiotics group (post-pre intervention).

| | Size<br>(# voxels) | MNI (x, y, z) | $p_{\text{FWE-cor}}$ at cluster level |
| --- | --- | --- | --- |
| R lat PFC | 967 | 40, 12, 20 | < 0.001 |
| R Caudate nucleus | 1275 | 6, 4, 8 | < 0.001 |
| R frontopolar cortex | 232 | 22, 64, 10 | 0.001 |
| pre-SMA | 558 | 4, 18, 54 | < 0.001 |
| L lat PFC | 468 | -56, 16, 22 | < 0.001 |

Note. lat PFC = lateral prefrontal cortex; pre-SMA = pre-supplementary motor area

**Figure S1.**
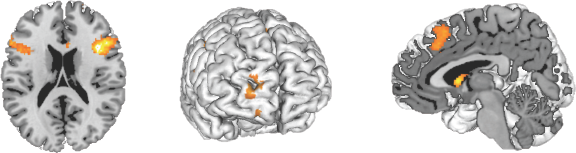
Negative correlation between stress-related DS backwards performance (post- minus pre-intervention) and incongruent versus congruent responses during the color-word Stroop task (post- minus pre-intervention) in the probiotics group. Only significant clusters are shown (whole-brain p_FWE_<.05, cluster defining threshold: p<.001).

